# A single domain shark antibody targeting the transferrin receptor 1 delivers a TrkB agonist antibody across the blood brain barrier to provide full neuroprotection in a mouse model of Parkinson’s Disease

**DOI:** 10.1101/2020.03.12.987313

**Authors:** Emily Clarke, Liz Sinclair, Edward J R Fletcher, Alicja Krawczun-Rygmaczewska, Susan Duty, Pawel Stocki, J Lynn Rutowski, Patrick Doherty, Frank S Walsh

## Abstract

Single domain shark antibodies that bind to the transferrin receptor 1 (TfR1) on brain endothelial cells can be used to shuttle antibodies and other cargos across the blood brain barrier (BBB). We have fused one of these (TXB4) to differing regions of TrkB and TrkC neurotrophin receptor agonist antibodies (AAb) and determined the effect on agonist activity, brain accumulation and engagement with neurons in the mouse brain following systemic administration. The TXB4-TrkB and TXB4-TrkC fusion proteins retain agonist activity at their respective neurotrophin receptors and in contrast to their parental AAb they rapidly accumulate in the brain reaching nM levels following a single IV injection. Following SC administration, the most active TrkB fusion protein, TXB4-TrkB_1_, associates with and activates ERK1/2 signalling in TrkB positive cells in the cortex and tyrosine hydroxylase (TH) positive dopaminergic neurons in the substantia nigra pars compacta (SNc). When tested in the 6-hydroxydopamine (6-OHDA) mouse model of Parkinson’s disease (PD) TXB4-TrkB_1_, but not the parental TrkB AAb or a TXB4-TrkC_1_ fusion protein, completely prevented the 6-OHDA induced death of TH positive neurons in the SNc. In conclusion, the fusion of the TXB4 TfR1 binding module allows a TrkB AAb to reach neuroprotective concentrations in the brain parenchyma following systemic administration.

## Introduction

The interaction of neurotrophins (NGF, BDNF, NT3 and NT4) with their cognate Trk receptors (TrkA, TrkB and TrkC, respectively) protects neurons from naturally occurring cell death during development ^1,2^. Their ability to nurture developing neurons spawned numerous studies to determine if they can promote the survival of adult neurons, particularly in the context of neurodegenerative disease or acute brain injury ^3,4^. In this context promising results have been found with BDNF which by activating the TrkB receptor can protect neurons from death in, for example, preclinical models of Parkinson’s disease (PD) ^5^, Alzheimer’s disease (AD) ^6,7^ and ischemic lesions ^8-11^. In addition, BDNF can promote functional recovery of injured neurons following spinal cord injury ^12-14^ and stimulate the production of new neurons in the adult brain ^15,16^. Loss of BDNF has also been suggested as a contributory factor to the progression of PD ^17-19^, AD ^20^ and Huntington’s disease (HD) ^21-23^, as well as conditions such as depression ^24,25^.

However, the therapeutic potential of BDNF in neurodegenerative diseases, acute brain injury and other neurological conditions has not been realised in the clinical setting due in part to a short plasma half-life *in vivo* ^26^, exclusion from the brain parenchyma following systemic administration, and poor diffusion throughout the parenchyma due to a high isoelectric point ^27^. The identification of agonist antibodies (AAb) that directly bind the TrkB receptor and mimic the neurotrophic activity of BDNF provides a clear pathway for the development of reagents with a long *in vivo* half-life, but the challenge of poor blood brain barrier (BBB) penetration remains. This has generally limited the use of the systemic delivery of TrkB AAb to peripheral disorders such as obesity ^28,29^ and peripheral neuropathy ^30^. Nonetheless, when delivered directly across the BBB by intracerebroventricular injection prior to an ischemic injury the TrkB AAb 29D7 enhances neuronal survival and promotes functional recovery ^31^.

There is considerable interest in the possibility of utilizing receptor mediated transcytosis pathways that exist on brain endothelial cells that form the BBB to carry biotherapeutics from the blood to the brain parenchyma with the transferrin receptor 1 (TfR1) being the most widely studied ^32^. TXB4 is a single domain shark variable new antigen receptor (VNAR) antibody specific to TfR1 and was developed from previously reported parental TXB2 VNAR ^33^ by restricted randomisation of CDR3 domain that led to improved brain penetration (JLR & PS, manuscript in preparation). TXB2 when fused to human IgG1 Fc (TXB2-hFc) readily crossed the intact BBB, reaching low nanomolar concentrations with a half-life of several days in the brain where it can be found to localised to TfR1 expressing neurons.

The 29D7 TrkB AAb mimics BDNF and has EC_50_ values of ∼50-200pM in a range of biological assays including neuronal survival ^34-36^. We hypothesised that if the TXB4 brain penetrating module was fused to this antibody it would accumulate in the brain following systemic administration at nM concentration sufficient to provide neuroprotection following disease or injury.

In the present study we have generated a novel TrkB AAb by cloning the variable regions of the mouse 29D7 TrkB AAb into a human IgG1 backbone that carries mutations in the Fc domain that attenuate effector function. This AAb has been further modified by fusion of the TXB4 module at the amino terminal domain of the IgG heavy chain (HC2N format) or between the CH1 and CH2 of the heavy chain (HV2N format). We have also designed a similar HC2N formatted TrkC AAb (TXB4-TrkC_1_) to use as a control in a limited set of experiments.

Our results show that unlike their parental antibodies the TXB4-TrkB and TXB4-TrkC_1_ fusion proteins rapidly accumulate in the brain reaching nM levels following a single IV injection. In the case of the TrkB AAb, the HV2N formatted version was ∼ 7-fold more active as a TrkB agonist than the HC2N formatted version in a cellular assay. The HV2N formatted version (TXB4-TrkB_1_) was taken forward for evaluation *in vivo*. We show that following SC administration TXB4-TrkB_1_ associates with and activates ERK1/2 signalling in TrkB positive cells in the cortex and tyrosine hydroxylase (TH) positive dopaminergic neurons in the substantia nigra compacta (SNc). When tested in the mouse 6-OHDA model of PD TXB4-TrkB_1_ completely prevented the loss of TH positive neurons throughout the SNc. In contrast, the parental TrkB AAb and TXB4-TrkC_1_ did not activate ERK signalling in TH positive neurons or protect them in the disease model. In conclusion, fusion with the TXB4 module allows the TrkB AAb to reach neuroprotective concentrations in the brain parenchyma following systemic administration generating a new class of biologic with therapeutic potential in a wide range of neurodegenerative diseases, acute brain injury situations and possibly depression.

## Materials and Methods

### Generation of the TrkB and TrkC Aab

Total RNA isolated from the mouse 29D7 hybridoma was subjected to whole transcriptome shotgun sequencing. The DNA and protein sequences of the mature primary VH and VL domains with their CDRs regions were identified using the Kabat numbering scheme. The 29D7 mouse VH and VL regions were cloned into constant regions of human heavy chain IgG1 and human light chain kappa, respectively. We have also used a TrkC AAb for a limited number of control experiments ^37^. The VH and VL regions of the monoclonal antibody 6.4.1, a potent and highly selective TrkC AAb ^38^ were cloned into constant regions of the human heavy chain IgG1 and human light chain kappa respectively. In all constructs, the human Fc domain contained the LALA double mutation (Leu234Ala together with Leu235Ala) to attenuate effector function^37^.

### Design and production of TXB4 fusion proteins

The brain uptake properties of TXB2 have been improved by restricted mutagenesis of the CDR3 region to generate TXB4 (JLR and PS, manuscript in preparation). For the TrkB AAb two TXB4 fusions were designed, one with the VNAR fused to the N-terminus of the TrkB heavy chain VH domain via 3xG4S linker (HC2N format) and one fused between the CH1 and CH2 domains via 3xG4S and 1xG4S linkers (HV2N format – TXB4-TrkB_1_), respectively. For the TrkC AAb, the same strategy was used to design a single HC2N formatted TXB4 construct (TXB4-TrkC_1_).

The proteins were expressed in CHO cells following transient transfections. The supernatant was collected and filtered using 0.22µm membrane filters and loaded onto HiTrap MabSelect SuRe Protein A (GE Healthcare) column pre-equilibrated against PBS, pH7.4. Protein A affinity bound proteins were eluted with 0.1M glycine, pH3.5 into neutralizing buffer (1M Tris-HCl pH9.0) and the buffer exchanged to PBS, pH7.4 using HiPrep 26/10 Desalting (GE Healthcare). Purity of the purified protein samples was determined by analytical size exclusion chromatography (SEC) using a Superdex200 column.

### TrkB and TrkC receptor reporter assay

TrkB and TrkC-NFAT-bla CHO-K1 reporter cell lines were purchased from Invitrogen. Cells were passaged twice a week in growth media consisting of DMEM-GlutaMAX medium (Gibco) supplemented with 10% dialysed fetal bovine serum, 100U/ml penicillin, 100µg/ml streptomycin, 5µg/ml blasticidin, 200µg/ml zeocin, 0.1mM non-essential amino acid solution (NEAA), and 25mM HEPES buffer. All additional reagents for tissue culture were purchased from Sigma. For the assay, 2 × 10^4^ cells were seeded in each well of a black-wall clear-bottom 96-well plate (Corning, USA) in 100μl of assay media consisting of DMEM-GlutaMax medium supplemented with 0.5% dialysed fetal bovine serum, 100U/ml penicillin, 100µg/ml streptomycin, 0.1mM NEAA and 25mM HEPES. Cells were incubated overnight at 37°C, 5% CO_2_. Parental antibodies and TXB4 fusion proteins were prepared in assay media and 50μl added per well to achieve a final concentration range as indicated in the results. Cells were incubated with the test reagents for 4 hr at 37°C, 5% CO_2_ before the manufacturer’s recommended assay protocol was followed. In brief, 30μL membrane-permeant fluorescent substrate coumarin cephalosporin fluorescein-acetoxymethyl ester (CCF4-AM and CCF2-AM, ThermoFisher) was added per well and incubated at room temperature for 90 min while protected from light. Since β-lactamase cleaves CCF2 substrate causing a shift in FRET emission, the excitation filter was set at 405nm, and the emission filters at 460 and 530nm on the plate reader (FlexStation, Molecular Devices, USA). The ratio of the two emission wavelengths (λ1/λ2) was calculated as a measure of β-lactamase activity, which in turn is an indicator of TrkB or TrkC receptor activation.

### Animal studies

All *in vivo* studies were performed in accordance with UK Animals Scientific Procedures Act (1986) and were approved by King’s College London Animal Welfare and Ethical Review Body. A total of 95 adult BalbC mice (8-12 weeks old, Envigo) were used: 75 for the brain accumulation study, 3 for the brain localisation studies and 17 for the 6-OHDA neuroprotection study. All animals were maintained on a 12:12 hour light:dark cycle with food and water available *ad libitum*

### Brain accumulation of TrkB AAb and TXB4 fusion constructs by ELISA

Female BalbC mice were injected IV with 3.6mg/kg (25nmol/kg) of either TrkB AAb (n=15), TrkC AAb (n=15) or with 4.3mg/kg (25nmol/kg) of TXB4-TrkB-HC2N (n=15), TXB4-TrkB-HV2N (n=15) or TXB4-TrkC-HC2N (n=15). Animals (n=3 per treatment group) were euthanized at 30 min, 1, 2, 4 or 18 hr post injection by overdose of phenobarbital (Euthatal; 200mg/ml, 1ml injection) before being intracardially perfused with phosphate-buffered saline (PBS) and the brain dissected into left and right hemispheres and stored at -80°C.

Brain accumulation was assessed by ELISA. Brains were homogenized in 3:1 (v/w) of PBS containing 1% Triton X-100 supplemented with protease inhibitors (cOmplete™, Sigma) using the TissueRuptor (Qiagen) at medium speed for 10 sec and then incubated for 30 min on ice. Lysates were spun down at 17,000 x g for 20 min, the supernatant was collected and blocked in 2.5% milk in PBS with 0.1% Tween 20 overnight at 4°C. MaxiSorp plates (Thermo) were coated with 100µl of goat anti-human Fc antibody (Sigma) diluted 1:500 in PBS overnight at 4°C. The plates were washed and incubated with blocking buffer for 1 hr at room temperature. Blocked brain lysates (100µl) were added to the blocked plates and incubated for 1 hr at room temperature. Plates were washed before detection and goat anti-human Fc-HRP conjugated antibody (Sigma) (1:5000 dilution in PBS) was added to the plate for a 1 hr incubation. Subsequently plates were washed and developed using TMB solution and 1% HCl was used to stop the reaction. The absorbance was measured at 450nm wavelength. The concentrations were determined using standard curves prepared for each construct individually.

### Brain localisation of TrkB AAb, TXB4-TrkB_1_ and TXB4-TrkC_1_ by immunohistochemistry

The TrkB AAb, TXB4-TrkB_1_ or TXB4-TrkC_1_ were injected at 10mg/kg SC into male BalbC mice. Animals were euthanized 18 hr later by overdose of phenobarbital (Euthatal; 200mg/ml, 1ml injection) before being intracardially perfused with PBS followed by 10% neutral buffered formalin (Sigma). Brains were removed and submerged in 10% neutral buffered formalin for 24 hr before being embedded in paraffin wax. Serial 7µm sagittal sections were obtained for immunofluorescent staining.

Sections were dewaxed (2 × 5 min in xylene, 4 × 2 min 100% IMS) and endogenous peroxidases quenched by immersion in 3% H_2_O_2_ for 10 min. Antigen retrieval was performed by boiling sections in 1mM citric acid pH6.0 for 10 min. Blocking solution (3% porcine serum albumin in 0.05M tris buffered saline (TBS) pH7.6) was applied for 90 min before sections were incubated at 4°C overnight in a humidified chamber with primary antibody. The TrkB and TrkC AAb and TXB4 fusion proteins were all detected with a single primary anti-human IgG antibody (Vector biotinylated goat anti-human IgG BA-3000 1:500). Abcam polyclonal chicken anti-TH ab76442 1:1500 was used to detect TH+ neurons in the SNc. CST polyclonal rabbit anti-pErk1/2 #9101 1:250 was used to detect activated ERK1/2. Abcam polyclonal rabbit anti-TrkB ab18987 was used to detect the endogenous TrkB receptor. Primary antibody was washed off for 2 × 5 min in 0.025% Triton-X100 in TBS before sections were incubated with fluorescent secondary antibodies at 1:500 for 90 min at room temperature. If a biotinylated primary antibody was used as in the case of Vector anti-human IgG BA-3000 sections were incubated with streptavidin-biotinylated HRP conjugate (Vectastain Elite ABC Kit, PK6100, VectorLabs) for 30 min at room temperature prior to addition of a streptavidin conjugated fluorescent secondary antibody. Fluorescent secondary antibodies used at 1:500 include: streptavidin conjugated Alexa Fluor 647, Alexa Fluor donkey anti rabbit 488, Alexa Fluor goat anti-chicken 488 and Alexa Fluor donkey anti-rabbit 647.

Secondary antibody was washed off sections for 2 × 5 min in 0.025% Triton-X100 in TBS before incubation with Sudan Black B 0.1% in 70% ethanol for 20 min at room temperature to quench autofluorescence. After rinsing off Sudan black under running water, slides were dried, and coverslips applied using Vectashield vibrance antifade with DAPI (Vector Labs H-1800). Fluorescence images were acquired using Zeiss 710 confocal microscope and Axiovision image analysis software.

### 6-OHDA unilateral lesion model of Parkinson’s disease

A single sub-cutaneous (SC) injection of either 5ml/kg PBS (n=5), or 5mg/kg of the TrkB AAb (n=4), TXB4-TrkB_1_ (n=4) or TXB4-TrkC_1_ (n=4) was given to male BalbC mice 24 hr prior to lesioning. A second injection of PBS or the above constructs at a dose of 2.5mg/kg was administered at post-lesion day 7.

For 6-OHDA lesioning, following anaesthesia induction (5% isofluorane/oxygen), animals were placed in a stereotaxic frame with blunt ear bars. Anaesthesia was maintained at 3% isofluorane/oxygen and body temperature maintained at 37 °C. The surgical site was sterilised with 0.4% chlorhexidine (Hibiscrub) before making an antero-proximal incision along the scalp. Fine-bore holes (Ø 0.5mm) were made in the skull at coordinates AP: +0.5mm and ML: +2.2mm (relative to bregma and skull surface) through which a blunt-ended 30-gauge needle was inserted to DV: −3.5mm. 6-OHDA.HBr (4μg in 3μl 0.02% ascorbate/saline) was infused unilaterally into the striatum (0.5μl/min) and the needle withdrawn 5 min later. This dose was predicted to produce a partial lesion over a 2-week period ^39^. After suturing, animals received a single dose of buprenorphine (Vetergesic; 0.1mg/kg; SC) for analgesia and 1ml of rehydrating Hartmann’s solution was administered SC daily for 5 days. One animal in the parental TrkB AAb group failed to recover adequately from surgery and was therefore excluded from the study.

### Immunohistochemical assessment of TH+ cell bodies in the SNc

On post-lesion day 14, animals were euthanized by overdose of phenobarbital (Euthatal; 200mg/ml, 1ml injection) before being intracardially perfused with PBS followed by 10% neutral buffered formalin (Sigma). Brains were removed and submerged in 10% neutral buffered formalin for 24 hr before being embedded in paraffin wax. Serial 7µm coronal sections encompassing the rostral, medial and caudal SNc were obtained and processed for TH staining.

Sections were dewaxed (2 × 5 min in xylene, 4 × 2min 100% IMS) and endogenous peroxidases quenched by immersion in 3% H_2_O_2_ for 10 min. Antigen retrieval was performed by boiling sections in 1mM citric acid pH6.0 for 10 min. Blocking solution (1% bovine serum albumin (BSA) in 0.05M tris buffered saline (TBS) pH7.6) was applied for 10 min before sections were incubated at room temperate overnight in a humidified chamber with primary polyclonal rabbit anti-TH antibody at 1:500 (ab152, Millipore). Primary antibody was washed off sections for 5 min in TBS before sections were incubated with biotinylated goat anti-rabbit secondary antibody at 1:500 at room temperature for 1 hour (BA-1000, VectorLabs). Secondary antibody was washed off sections for 5 min in TBS before incubation with streptavidin-biotinylated horseradish peroxidase conjugate (Vectastain Elite ABC Kit, PK6100, VectorLabs) for 30 min at room temperature. Colour was developed using 3-3’-diaminobenzidine (DAB) substrate kit (DAB Peroxidase HRP Substrate Kit, SK-4100, VectorLabs). Finally, sections were rinsed thoroughly in distilled H_2_O for 10 min to remove any trace of DAB and then dehydrated in 100% IMS (4 × 2 min) and cleared in xylene (2 × 5 min). Slides were mounted with coverslips using the solvent based plastic mountant DPX (Sigma).

Photomicrographs of TH-immunostained SNc sections (n = 3-6 sections per mouse at each of the caudal [AP: −3.52mm], medial [AP: −3.16mm] and rostral [AP: −2.92mm] levels relative to bregma) were acquired at 20× magnification using a Zeiss Apotome microscope and Axiovision software (Carl Zeiss Ltd). ImageJ software was used to manually count viable (i.e. intact round cells with a clear nucleus and cytoplasm) TH-positive A9 dopaminergic cells of the SNc in both the lesioned and intact hemispheres. SNc cell number in the lesion hemisphere was calculated as percentage of that lost in the intact hemisphere. Data were combined across all three rostro-caudal levels to generate a single average value for each animal. Mean data were then calculated per treatment group.

### Statistical analysis

All statistical analysis was performed using Graphpad Prism 8 software. The concentration response curve of TrkB constructs in the NFAT reporter assay was analysed by non-linear regression to give EC_50_ values (n=3-5 independent experiments per concentration per treatment). Percentage TH+ cell loss in the lesioned relative to the intact SNc in the 6-OHDA PD mouse model was tested for Gaussian distribution by Shapiro-Wilk and parametric statistics applied accordingly. The percentage TH+ cell loss of the lesioned relative to intact SNc was compared across treatment groups by one-way ANOVA followed by Tukey’s HSD, *p=<0.05, n=3-5 animals per group.

## Results

### Activity of the TrkB AAb and TXB4-fusion proteins in a reporter assay

The TrkB-NFAT-bla CHO-K1 cell line expresses the human TrkB receptor and provides a simple quantitative assay for receptor activation ^36^. The specificity of 29D7 for TrkB and its activity in the NFAT TrkB reporter assay relative to BDNF has been comprehensively described by others ^34,35^. In the present study we tested the modified 29D7 (TrkB AAb) and the two TXB4 fusion proteins in the same assay, presenting their responses as a percentage of the response maximally achieved by TrkB AAb (Fig. 1).

**Fig1.**
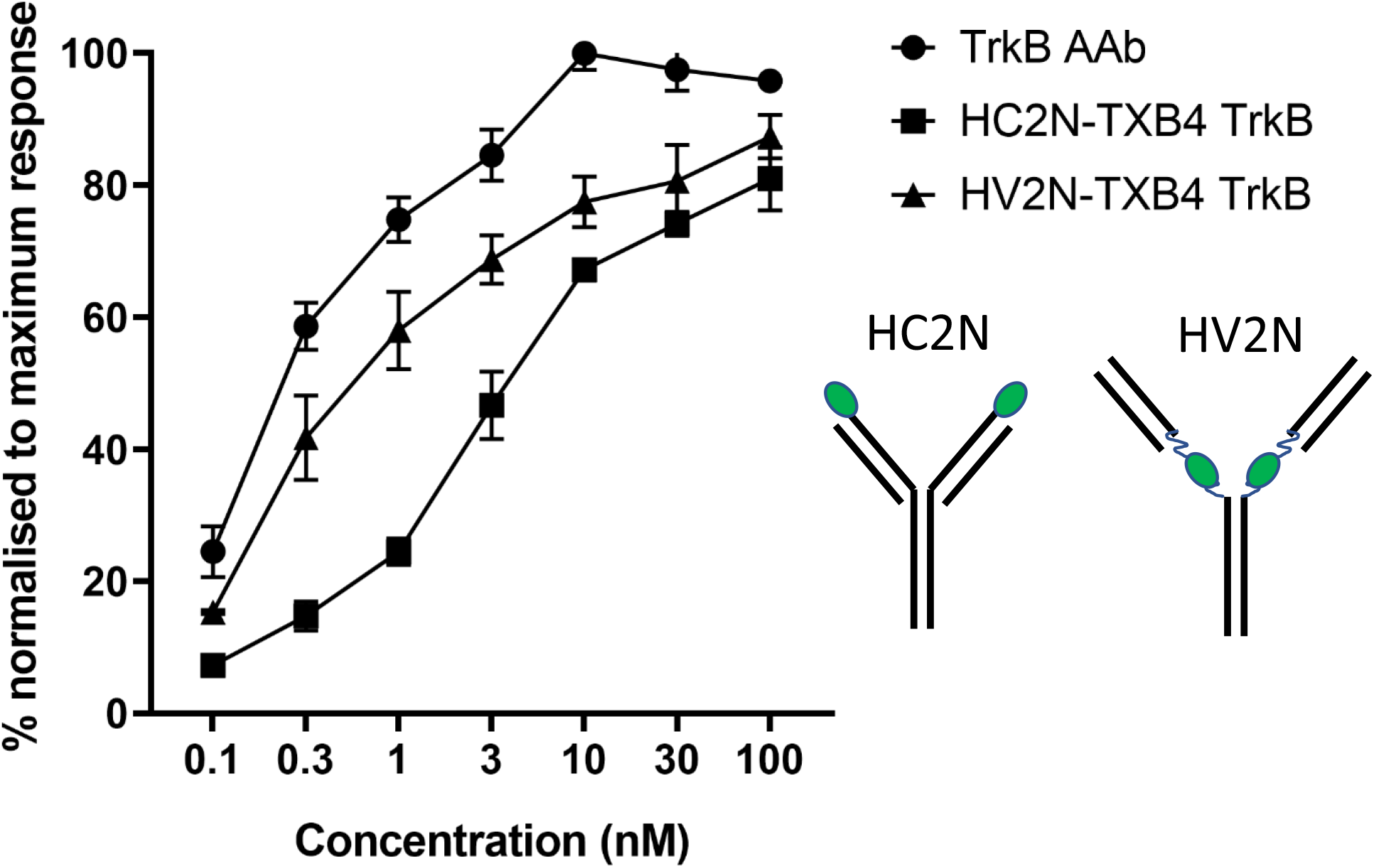
Agonist activity of TrkB AAb and TXB4-fusion proteins. The parental TrkB AAb and two TXB4 fusion proteins (as illustrated in cartoon with the green module representing the TXB4 module) were tested for agonist activity in the TrkB-NFAT-bla CHO-K1 reporter cell line assay (see methods for details). Data are presented as the mean ± S.E.M pooled from 3-5 independent experiments and are normalised to the maximal response elicited by the parental TrkB AAb in each experiment

With the TrkB AAb a significant activation of the TrkB receptor was clear at 100pM with an EC_50_ of 0.53nM determined from several independent experiments. The dose response curves for both TXB4 fusion proteins were shifted to the right, but both reagents reached the same maximal response as the parental TrkB AAb. Interestingly, the HV2N formatted version (EC_50_ = 0.75nM) was approximately 7-fold more potent compared with the HC2N format (EC_50_ = 5.18nM) with substantial activity (∼40% of the maximum response) seen at 300pM. In a parallel set of experiments the agonist activity of TXB4-TrkC_1_ was evaluated in the TrkC-NFAT-bla CHO-K1 assay. TXB4-TrkC_1_ was a full agonist with an EC_50_ of 1.6nM.

### Brain accumulation of the TrkB and TrkC AAb and the TXB4 fusion proteins

Mice were injected IV with 25nmol/kg of either the TrkB or TrkC AAb or the above TXB4 fusion proteins with plasma concentration and brain accumulation determined by ELISA assay after 0.5, 1, 2, 4 and 18 hr (see methods for details). For the TrkB proteins plasma measurements were not significantly different between groups at the 30 min time point (ranging from 350-450nM) or at the 18-hr time point (ranging from 100-200nM) (data not shown). However, brain accumulation differed markedly. Where the parental TrkB AAb showed very little accumulation over the 18-hr period with a measured value of 0.39 ± 0.08nM (Fig 2) a time-dependent accumulation of both TXB4 fusion proteins was seen with increases in the brain concentration relative to the parental antibody apparent at the first measured timepoint (30 min) with a steady accumulation seen over the 18-hr period resulting in concentrations of 4.69 ± 0.34 and 11.52 ± 0.40 nM for the HV2N and HC2N formats respectively (all values mean ± S.E.M, n=3 animals). Despite the observation that the HC2N format accumulates in the brain at higher levels than the HV2N format (aka TXB4-TrkB_1_), we took the later forward for further study based on the 7-fold higher agonist activity measured in the TrkB reporter assay (Fig 1). In an independent set of experiments (using same methods) we measured brain accumulation of the parental TrkC AAb and TXB4-TrkC_1_; at the 18 hr timepoint the parental TrkC AAb value was 0.33 ± 0.10nM while TXB4-TrkC_1_ was 6.47 ± 0.57nM (mean ± S.E.M, n=3 animals).

**Fig2.**
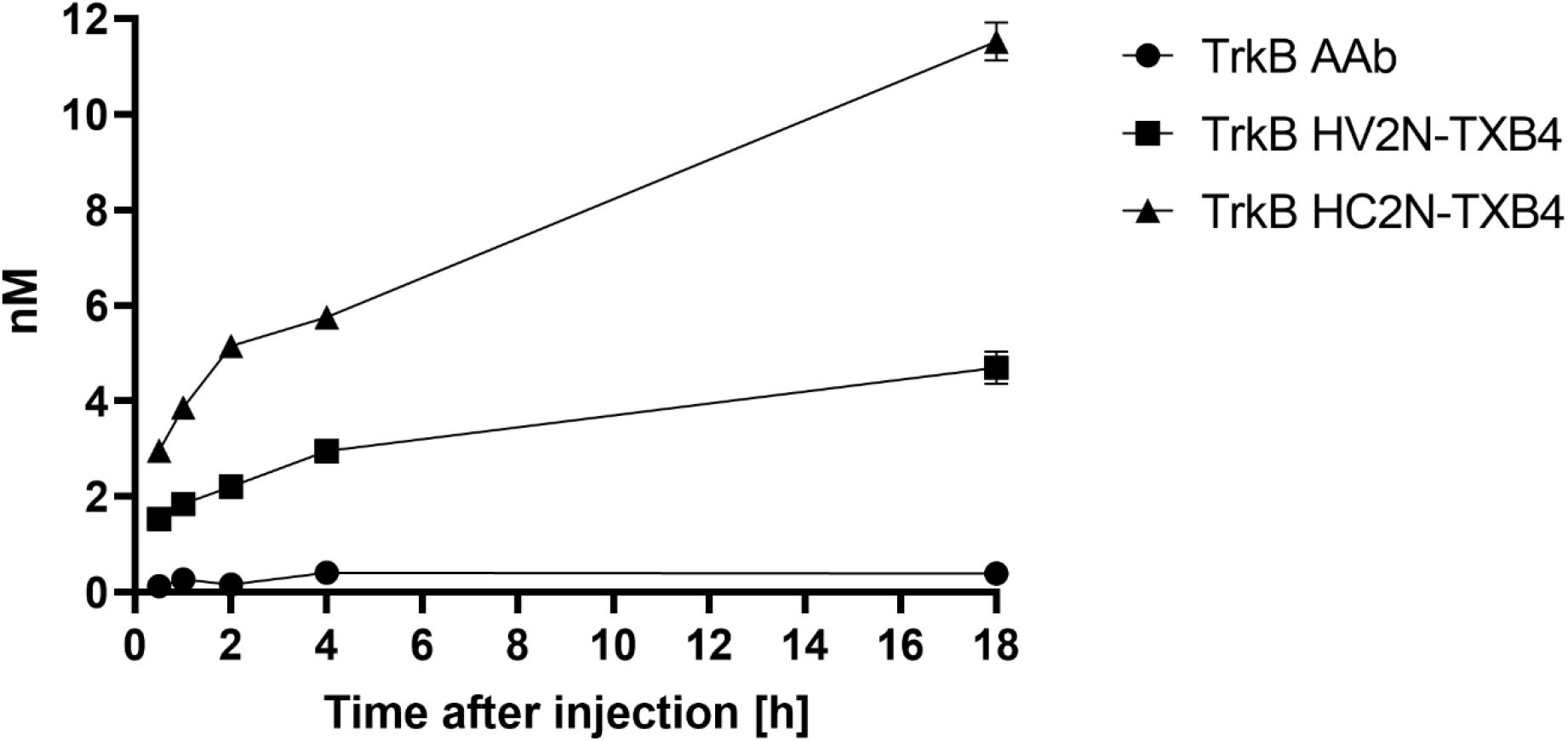
Brain uptake of TrkB AAb and TXB4 fusion proteins. The TrkB AAb and TXB4 fusion proteins were administered by single IV injection. Brains were perfused and harvested and concentration of the three agonist antibodies determined at the given timepoints (see methods for details). Data are presented as the mean ± S.E.M from three independent animals at each time point for each treatment.

### Brain localisation of TrkB AAb and TXB4-TrkB_1_

We next used an anti-human IgG antibody to determine if we could see where the TrkB antibodies localise in the brain. To this end the parental TrkB AAb or TXB4-TrkB_1_ were administered by a single SC injection (10mg/kg) and 18 hr later the perfused brains were isolated and processed for immunohistochemistry as described in the methods. At lower magnification we fail to see any TrkB AAb staining anywhere in the brain (e.g. see Fig 3a). This is in stark contrast to our ability to readily detect TXB4-TrkB_1_ within endothelial cells that line capillaries throughout the brain (e.g. see Fig 3b). If the TrkB AAb or TXB4-TrkB_1_ engage TrkB receptors in cells we would expect them to be internalised into those cells as this is a canonical feature of target engagement ^40^. The most conspicuous staining for the TrkB receptor was an intracellular staining of large neurons in the cortex (inserts in Fig 3c and Fig 3d). We did not detect any TrkB AAb in these cells even at high magnification (Fig 3c), but in contrast we could readily detect TXB4-TrkB_1_ in these cells (Fig 3d). Activation of the ERK1/2 signalling pathway is a canonical feature of neurotrophin receptor activation, and this can be detected by antibodies that recognise the phosphorylated form of ERK1/2 ^2^. Activated ERK1/2 was not detected by this method in the brains of animals treated with the TrkB AAb (e.g. see Fig 3e), however it was readily detected in the brain following treatment with TXB4-TrkB_1_ and this was most obvious in the cortex (Fig 3f).

**Fig3.**
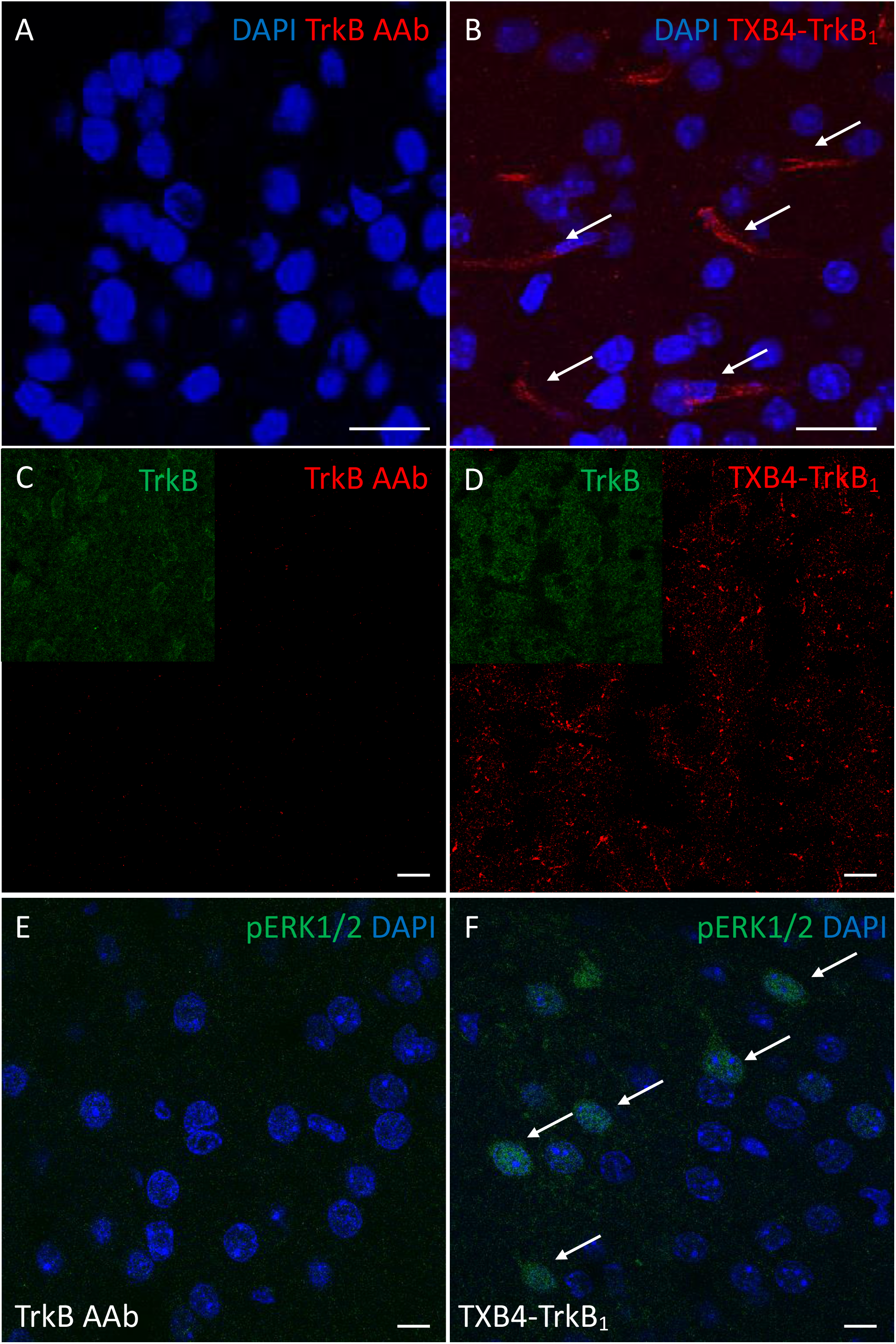
Localisation of human IgG and pERK1/2 in brains of animals treated with the TrkB AAb or TXB4-TrkB_1_. The TrkB AAb or TXB4-TrkB_1_ were administered SC to mice (10mg/kg) and brains harvested after 18 hr and stained for human IgG (see methods for details). At x40, human IgG was not detected following TrkB AAb administration (A) but was readily seen in endothelial cells following TXB4-TrkB_1_ administration (B) with arrows highlighting some of the positive staining). At higher magnification (x63 oil immersion objective) human IgG was not detected in the cortex following TrkB AAb administration (C) but was readily seen as a punctate intracellular stain in large neurons following TXB4-TrkB_1_ administration (D). The inserts in both C and D show the expression of the TrkB receptor in the same field. The same cortical regions as in C&D were also stained with an antibody that recognises pERK1/2 (see methods for details). pERK1/2 was not detected at high magnification (x63) following TrkB AAb administration (E) but was readily detected in cells and nuclei (as indicated by arrows) following TXB4-TrkB_1_ administration (F). DAPI was used to label nuclei in A&B and E&F. The scale bar in all images = 10μm.

### Target engagement and neuroprotection in the 6-OHDA model of PD

When delivered locally BDNF limits the death of TH+ dopaminergic neurons in the SNc that normally accompanies the injection of 6-OHDA into the striatum ^41-44^; to this end we have tested if the systemic administration of TXB4-TrkB_1_ is neuroprotective in this model of PD. We first wanted to determine if the systemically administered TXB4-TrkB_1_ localises to the SNc in a healthy animal, using tissue from the above-mentioned brain localisation studies. As expected, we did not detect the presence of the TrkB AAb within the vicinity of TH+ cells in the SNc 18 hr after SC delivery (Fig 4a). In contrast, TXB4-TrkB_1_ can readily be detected in this area of the brain 18 hr after SC administration (Fig 4b; 10mg/kg in both cases). Moreover, while ERK1/2 is not activated in TH+ neurons following TrkB AAb administration (Fig 4c), it is clearly activated in several TH+ neurons following TXB4-TrkB_1_ administration (Fig 4d). In contrast to TXB4-TrkB_1_ we were unable to detect the TXB4-TrkC_1_ in the SNc at 18 hr following a 10mg/kg SC injection and did not detect pERK1/2 in TH+ cells (data not shown).

**Fig4.**
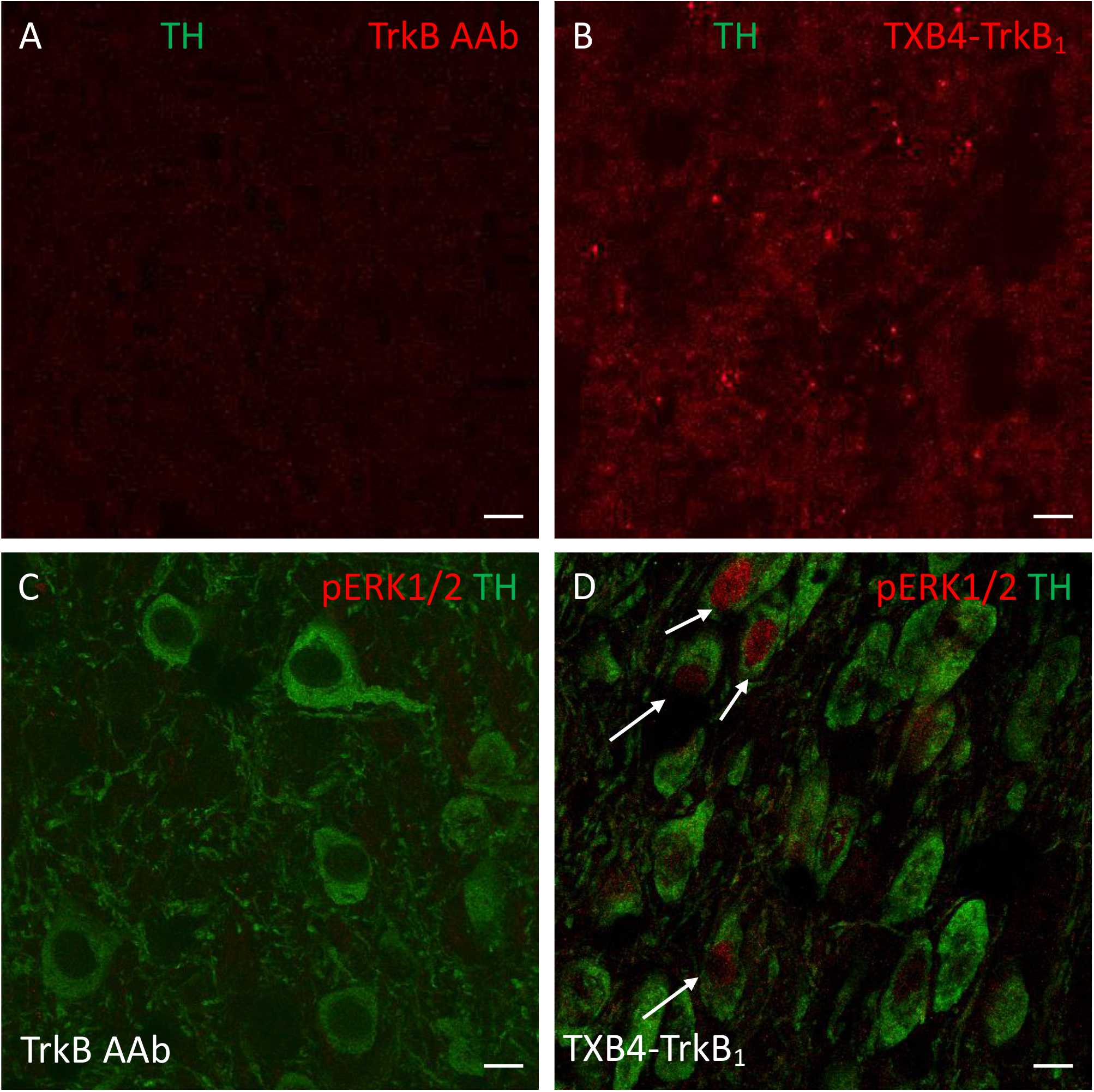
Localisation of human IgG and pERK1/2 in the SNc of animals treated with TrkB AAb or TXB4-TrkB_1_. TrkB AAb or TXB4-TrkB_1_ were administered SC to mice (10mg/kg) and brains harvested after 18 hr and stained for human IgG in green or pERK1/2 in red (see methods for details). Within the SNc human IgG was not detected as a consequence of TrkB AAb administration (A) but was readily seen in endothelial cells following TXB4-TrkB_1_ administration (B). The inserts in both show dopaminergic neurons within the same field revealed by co-staining for TH. The same regions as in A&B were also co-stained for TH and pERK1/2 (see methods for details). pERK1/2 was not detected at high magnification (x63) following TrkB AAb administration (C) but was readily detected in some cell nuclei (as indicated by arrows) following TXB4-TrkB_1_ administration (D). Scales bars = 10µm.

To test for neuroprotection, mice were treated with either PBS (control), TXB4-TrkB_1_, TrkB AAb or TXB4-TrkC_1_ (5mg/kg) 24hr before inducing a partial 6-OHDA lesion and again at day 7 (with reduced construct dose of 2.5mg/kg). 14 days after injection of 6-OHDA, brains were isolated and processed for TH immunoreactivity in the rostral, medial and caudal SNc as described in material and methods. As expected, there was a clear reduction (27.30 ± 7.31%) in the number of TH+ cells as a percentage of the intact hemisphere following 6-OHDA treatment in the control group (Fig 5). In contrast there was essentially no cell loss (3.18 ± 2.6%) in TXB4-TrkB_1_ treated mice and this value was significantly less than that seen in the control group (One-way ANOVA, Tukey’s HSD, *p=0.035). Neuronal loss was still apparent and not significantly different from the control group with either the TrkB AAb (12.47 ± 2.97%) or TXB4-TrkC_1_ (20.74 ±4.11%) treatment (Fig 5) (all values mean ± S.E.M).

**Fig5.**
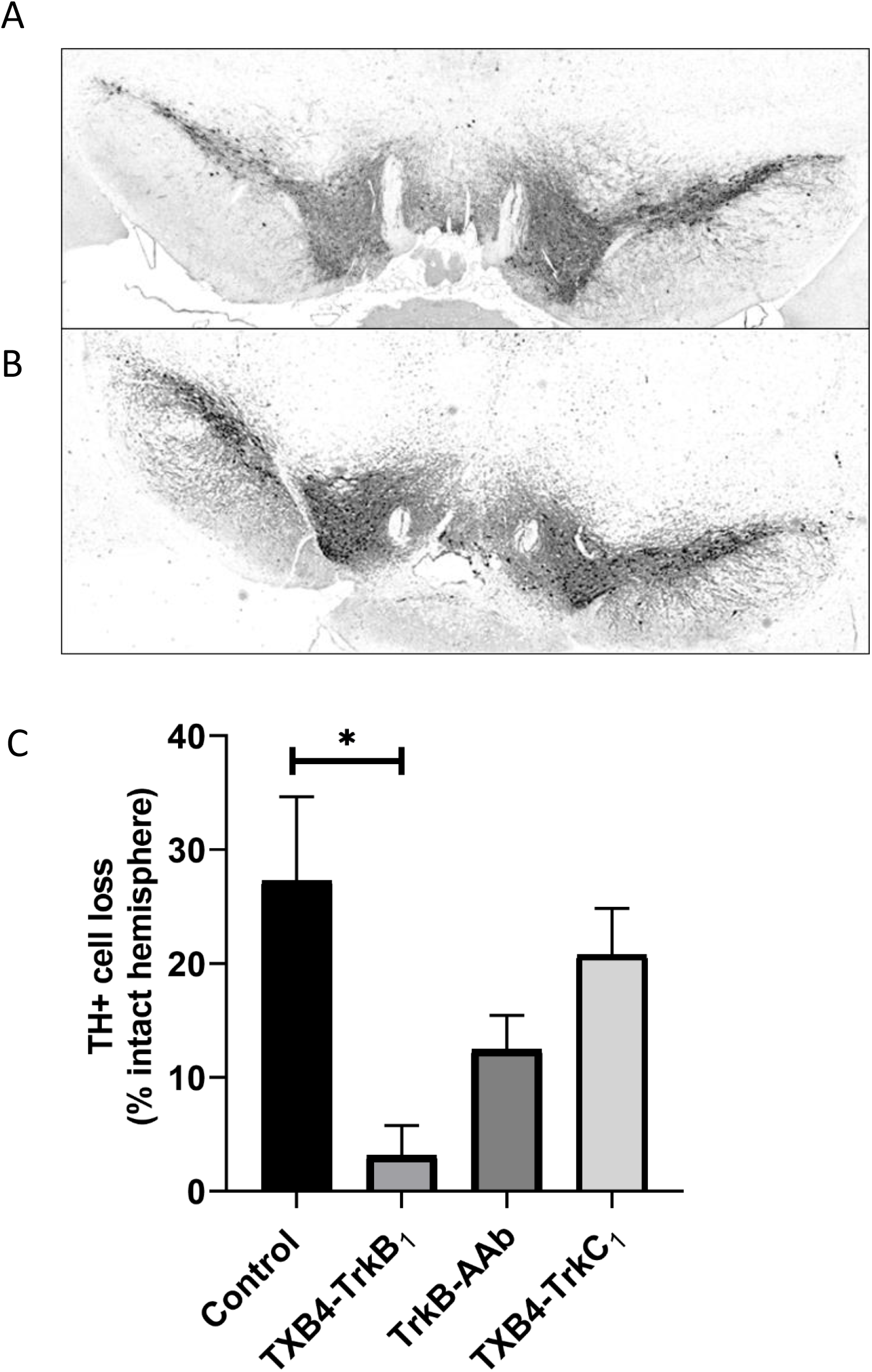
TXB4-TrkB_1_ protects dopaminergic cell bodies in the substantia nigra (SNc) against neuronal loss in the 6-OHDA mouse model of Parkinson’s disease. Representative images of TH+ dopaminergic SNc cells within the 6-OHDA lesioned (left) and intact (right) sides of PBS control (A) and TXB4-TrkB_1_ treated mice (B) x20 magnification, scale bar = 500μm. In (C) cell loss within the 6-OHDA lesioned (left) relative to control (right) side of the hemisphere is shown for the PBS control group and groups treated with TXB4-TrkB_1_, TrkB AAb or TXB4-TrkC_1_ as indicated (see methods for details). Data represented as mean± S.E.M % TH+ cell loss relative to intact hemisphere ± S.E.M. One-way ANOVA, Tukey’s HSD, *p=0.035, n=3-5 mice per treatment group.

## Discussion

The BDNF-TrkB signalling pathway is considered a drug target for a wide range of neurological diseases and depression (see introduction). However, a short half-life of approximately 10 min in plasma ^26^ and 1 hour in CSF ^45^, and a high isoelectric point (pI∼10) that limits its diffusion in tissues ^27^ have in part hampered clinical development. BDNF can also bind to the p75 neurotrophin receptor (p75^NTR^) and in some instances activation of this receptor can induce apoptosis ^46^. p75^NTR^ is also an integral component of a receptor complex that inhibits axonal growth and the interaction of BDNF with this complex might detract from its regenerative function ^47^. The BDNF/ p75^NTR^ interaction might also limit the therapeutic potential of BDNF.

TrkB antibodies do not bind p75^NTR^ and in general have a plasma half-life of several days. In terms of efficacy, the intravitreal delivery of the 29D7 TrkB AAb delays retinal ganglion cell death in models of acute and chronic retinal injury ^48,49^ whilst intracerebroventricular administration prior to initiation of a neonatal hypoxic-ischemic brain injury in rats significantly increased neuronal survival and behavioural recovery ^31^. Intrathecal application of 29D7 improves motor neuron survival and regeneration in models of spinal cord injury and motor neuron degeneration ^14^. It follows that there might be translational opportunities in the development of a version of 29D7 (or indeed another TrkB AAb) that can readily cross the BBB following systemic administration. The strategy of using TfR1 receptor antibodies to deliver cargos across the BBB is not novel ^50-52^. However, limitations include the retention of TfR1 antibodies within brain capillaries ^53,54^, the lysis of TfR1-expressing reticulocytes ^55^ and iron deficiency toxicities due to competition for transferrin binding to the TfR1 and/or antibody induced targeting of TfR1 to lysosomes ^56-59^. These are hopefully not insurmountable problems and interestingly cell type specific binding of a range of TfR1 antibodies, perhaps due to differential glycosylation, has been reported providing a rational for development of antibodies with unique tissue binding profiles ^60^.

The shark VNAR is a small 12-15 kDa compact domain with an extended CDR3 loop that can engage with cryptic epitopes inaccessible to standard immunoglobulins ^61^. The TfR1 binding TXB2 VNAR can carry cargos, including a human IgG Fc domain (TXB2-hFc) as well as biologically active neurotensin, across the BBB. Importantly the TXB2-hFc does not compete for transferrin binding, clear the TfR1 from capillaries, or lyse reticulocytes ^33^. TXB4 is a version of TXB2 with some modifications in the CDR3 that improved brain penetration (JLR and PS, manuscript in preparation). It shares the same binding epitope and consequently also does not compete for transferrin binding.

We have designed and synthesised two TXB4-TrkB AAb fusion proteins, one with an exposed VNAR domain at the N-terminus of the HC (HC2N), the other positioned between CH1 and CH2 domains above the hinge region (HV2N). One showed an ∼12-fold and the other a ∼30-fold increase in brain accumulation over an 18-hr period relative to the parental antibody, but this might be an underestimate as it is not clear if the parental antibody value of 0.39nM reflects genuine accumulation or non-specific binding as we cannot detect it in the brain by immunohistochemistry (see below). Interestingly whereas the HC2N configuration appears to favour brain accumulation the HV2N configuration appears to favour preservation of TrkB agonist activity. Conclusions cannot be drawn from such a limited study, but a trade-off between TrkB agonist activity and TfR1 dependent brain accumulation might be expected based on positioning the VNAR relative to the TrkB binding site. Nonetheless, this is a clear demonstration of the adaptable modular nature of the VNAR.

The HV2N formatted construct (which we have designated as TXB4-TrkB_1_) was approximately 7-fold more active in the TrkB reporter assay than the HC2N format with substantial activity (∼40% of the maximum response) seen at 300pM. It reaches a “concentration” of ∼4.7nM at 18 hr following a single IV injection of 25nmol and if as little as 10% of this is free in the brain parenchyma one would expect robust activation of the TrkB receptor. Target engagement and activation would be manifested by the uptake of TXB4-TrkB_1_ into TrkB expressing cells and activation of canonical signalling pathways such as the ERK1/2 cascade ^62^. Indeed, following a single SC injection in a healthy animal, we could readily detect TXB4-TrkB_1_ within the large TrkB positive neurons in the cortex and also found activation of the ERK1/2 cascade throughout the cortex as indexed by binding of an antibody that recognises the phosphorylated activated enzymes. Likewise, TXB4-TrkB_1_ was found in the SNc and activated ERK1/2 was clearly activated in the TH+ dopaminergic neurons in the SNc in the same animal. As a control, antibody accumulation and ERK1/2 activation were not seen in the SNc following systemic administration of the parental TrkB AAb.

When injected into the striatum 6-OHDA is taken up into dopaminergic neurons via the high-affinity dopamine transporter. It is then oxidised, and the toxicity of the released reactive oxygen species is reflected in a loss of dopaminergic tracts and terminals in the striatum and cell bodies in the SNc ^63^. This is a well-established preclinical model of PD ^64^, but it is noteworthy that toxicity due to generation of reactive oxygen species might be causative of neuronal loss in a wide range of other neurodegenerative diseases. In this study the unilateral administration of 6-OHDA was associated with the loss of 20% -30% of the neurons throughout the SNc. Remarkably, there was no significant neuronal loss throughout the SNc in animals treated with TXB4-TrkB_1_. Interestingly TXB4-TrkC_1_ also accumulated in the brain but did not localise to or activate ERK1/2 in the TH+ dopaminergic neurons nor offer any neuroprotection against the 6-OHDA lesion. The parsimonious explanation is that the localisation within the parenchyma and efficacy of TXB4-TrkB_1_ is driven by the TrkB and not the TfR1 binding activity, but future studies will have to test some aspects of this perhaps by the co-administration of selective TrkB antagonists alongside TXB4-TrkB_1_. Likewise, future studies will be required to determine the impact of TXB4-TrkB_1_ on the development of the motor deficits that are a well-studied trait in this model.

Antibodies can show efficacy in neurodegenerative models when administered without a shuttle due to passive transfer and/or uptake through a disease related or injury induced “leaky” BBB. Indeed, there is evidence suggesting the BBB is damaged to some extent for a limited period in the 6-OHDA models^65^. In this study the parental TrkB AAb showed a trend towards limited neuroprotection in the SNc, although it did not reach statistical significance. Thus, there might have been partial neuroprotection because of some leakage of the TrkB AAb across the compromised BBB, but importantly the TXB4-TrkB_1_ was by far the more effective treatment. Nonetheless, Han et al., has demonstrated the therapeutic potential of IV administered TrkB AAb Ab4B19 for ischaemic brain injury based on leakage across the compromised BBB at the site of injury ^66^. Indeed, many neurodegenerative diseases are associated with a comprised BBB related to neuroinflammatory processes ^67,68^ and this might in part explain the efficacy of, for example, the amyloid-β antibody bapineuzumab in AD ^69^. However, one needs to consider the likely beneficial effects of using a shuttle to deliver a therapeutic antibody to the brain. The most obvious of these is the fact that the therapeutic antibody can be delivered to regions of the brain where the BBB has yet to be compromised or indeed has recovered from damage. Likewise, the use of a shuttle is always likely to result in more antibody reaching the parenchyma increasing the probability of a therapeutic response. Finally, getting more antibody from the serum to the brain will obviously impact on the systemic dose required for a therapeutic effect. In this context it is noteworthy that in patients treated with the therapeutic anti-Aβ antibody aducanumab the number of individuals experiencing side effects such as microhemorrhages of the BBB and oedema increased from 13% to 47% when the administered dose went from 3 to 10mg/kg^70^.

In summary, this paper has shown for the first time that the TXB4 VNAR targeting the TfR1 can be fused to an AAb to allow efficient transport across the BBB and access to the brain parenchyma at an active concentration. In this regard TXB4-TrkB and TXB4-TrkC AAb crossed the BBB and accumulated in the brain at low nM concentrations. One of these (TXB4-TrkB_1_) engaged with and activated neurotrophin signalling in target cells that included neurons that are susceptible to loss in AD (cortical neurons) and PD (dopaminergic neurons in the SNc). Furthermore, systemic treatment with TXB4-TrkB_1_ prevented the neuronal loss normally seen in a partial lesion mouse model of PD. As such TXB4-TrkB_1_ can be considered as the first in class of a new generation of BBB penetrant AAb with therapeutic potential in a wide range of neurodegenerative diseases, acute brain and spinal cord injury situations and possibly depression.

## Nonstandard abbreviations

AAb: *agonist antibody*
AD: *Alzheimer’s disease*
BBB: *blood brain barrier*
CDR3: *complementarity-determining region 3*
HD: *Huntington’s disease*
PD: *Parkinson’s disease*
TfR1: *transferrin receptor 1*
TH: *tyrosine hydroxylase*
SNc: *substantia nigra pars compacta*
VNAR: *variable domain of new antigen receptor*.

